# Bet hedging in a unicellular microalga

**DOI:** 10.1101/2023.09.08.556835

**Authors:** Si Tang, Yaqing Liu, Katrin Hammerschmidt, Jianming Zhu, Xueyu Cheng, Lu Liu, Jin Zhou, Zhonghua Cai

## Abstract

Understanding how organisms adapt to unpredictable future environments is a fundamental goal in biology, which becomes even more urgent in an era of rapid climate change. One evolutionary adaptation to randomly fluctuating environments is bet hedging, a strategy that successfully facilitates reproduction and population persistence and has been widely reported from microbes to humans. Empirical evidence for its presence in microalga, one of Earth’s most important primary producers and carbon sinks, is lacking. Here, we report a bet-hedging strategy in the unicellular microalga *Haematococcus pluvialis.* In a series of experiments, we show that an isogenic *H. pluvialis* population reversibly diversifies into hetero-phenotypic mobile and non-mobile subunits, independent of environmental conditions. Mobile cells grow faster but are more susceptible to external stressors, while non-mobile cells hardly grow but are more stress-resistant. This is attributed to dramatic shifts from growth-promoting activities (cell division, photosynthesis) to resilience-promoting cellular metabolic processes, including cell enlargement and aggregation, and accumulation of antioxidant and energy-storaging compounds. Our results provide experimental evidence for bet hedging in microalga, which has implications for their potential to adapt to current and predicted future conditions, and thus for the conservation of ecosystem functions.

## Introduction

Ever since the ancient Earth, organisms needed to adapt to changing global environments. While most populations can accommodate regular environmental changes, such as daily diurnal and annual seasonal cycles, dealing with random changes or fluctuations poses a challenge. As far as our present-day Earth is concerned, the environment is changing at an unprecedented rate - mainly due to anthropogenic activities - and so generating novel and unexpected environmental fluctuations that pose urgent challenges to every living organism^1,2^. Therefore, the question of whether and how organisms adapt to unpredictable environments is of crucial importance.

In evolutionary theory, bet hedging, is proposed as a successful adaptive strategy to cope with randomly fluctuating environments^3,4^. Bet hedging in unicellular organisms means that an isogenic population can increase its long-term fitness at the expense of lower short-term fitness through the stochastic development of different phenotypes^5–8^. Different phenotypes may be well-adapted to different environmental conditions, and individuals with different phenotypes have varying reproductive success and survival rates that change depending on external conditions. Unlike real-time sensing and then responding to changes, bet hedging is advantageous, as it allows a subpopulation to display the phenotype that will be adaptive in a future environment before the environment shifts. Upon sudden environmental changes, such a bet-hedging strategy represents a rapid and reliable mechanism for avoiding extinction^4^.

Bet hedging has attracted a growing interest and has been reported in diverse research areas^9–16^, both theoretically and experimentally, and more and more evidence is emerging from various model organisms. To our knowledge, however, there has been no direct evidence for bet hedging in microalga, which are among the most important primary producers on Earth and are of critical ecological importance, participating in global biogeochemical carbon and nitrogen cycles^17–21^. As their “physiology and persistence are severely affected by global climate change”^22^, we urgently need to increase our understanding on how they adapt to fast-changing environments.

Here, the genetically clonal microalgal species *Haematococcus pluvialis* is used as a model organism to study bet hedging in microalga. *H. pluvialis* (Chlorophyceae, Volvocales) is a widely distributed, buoyant unicellular biflagellate freshwater microalga. Thanks to the commercial value of the strain, known as the best natural source for the bio-product astaxanthin, its morphological and physiological changes as well as metabolic variations have been widely studied under different cultural environments^18–20^. A particularly interesting phenomenon that has been repeatedly reported^23–26^ and that we have also observed in both culture flasks and bioreactors is the co-existence of two behaviorally distinct subpopulations, namely mobile and non-mobile green cells.

So far, it has been assumed that external stress factors lead to the transformation of mobile to non-mobile green cells, which turn into dormant red cells when conditions deteriorate further^27^. However, why mobile and non-mobile subpopulations coexist in the green stage remains unexplored, and the functional and ecological significance of that phenotypic heterogeneity is not understood.

As non-mobile cells occur much earlier than expected under standard culture conditions, we doubt that external stress factors are the true or only trigger for the observed phenotypic heterogeneity. Instead, we propose here that the co-existence of two phenotypes in the *H. pluvialis* populations can be explained by bet hedging (see Fig. 1A for a comparison of the stress-induced cellular transformation with the bet hedging hypothesis tested in this article). In the case of bet hedging, we expect the mobile subgroup of the population to produce more offspring but be more susceptible to external stressors, while the non-mobile subgroup should be more stress-resistant at the expense of lower reproduction. Therefore, in this study, we used laboratory populations of *H. pluvialis* to test the following: (1) Do the two subpopulations of *H. pluvialis* behave differently under standard and stress conditions? This would be expected if they have different stress resistance; (2) Does a hetero-phenotypic population show lower variance in reproductive fitness under varying environments as compared to a phenotypically uniform population? which is a canonical feature of bet- hedging populations^28,29^. To this end, we combined growth and survival experiments with microscopic, physiological and transcriptomic analyses. If our hypothesis is correct, this study provides the first experimental report of a bet-hedging behaviour in the phyla of microalga. This would offer fundamental insights into how these organisms can cope with unpredictable fluctuating environments and give a different perspective on future species or ecosystem rescue or management.

**Figure. 1.**
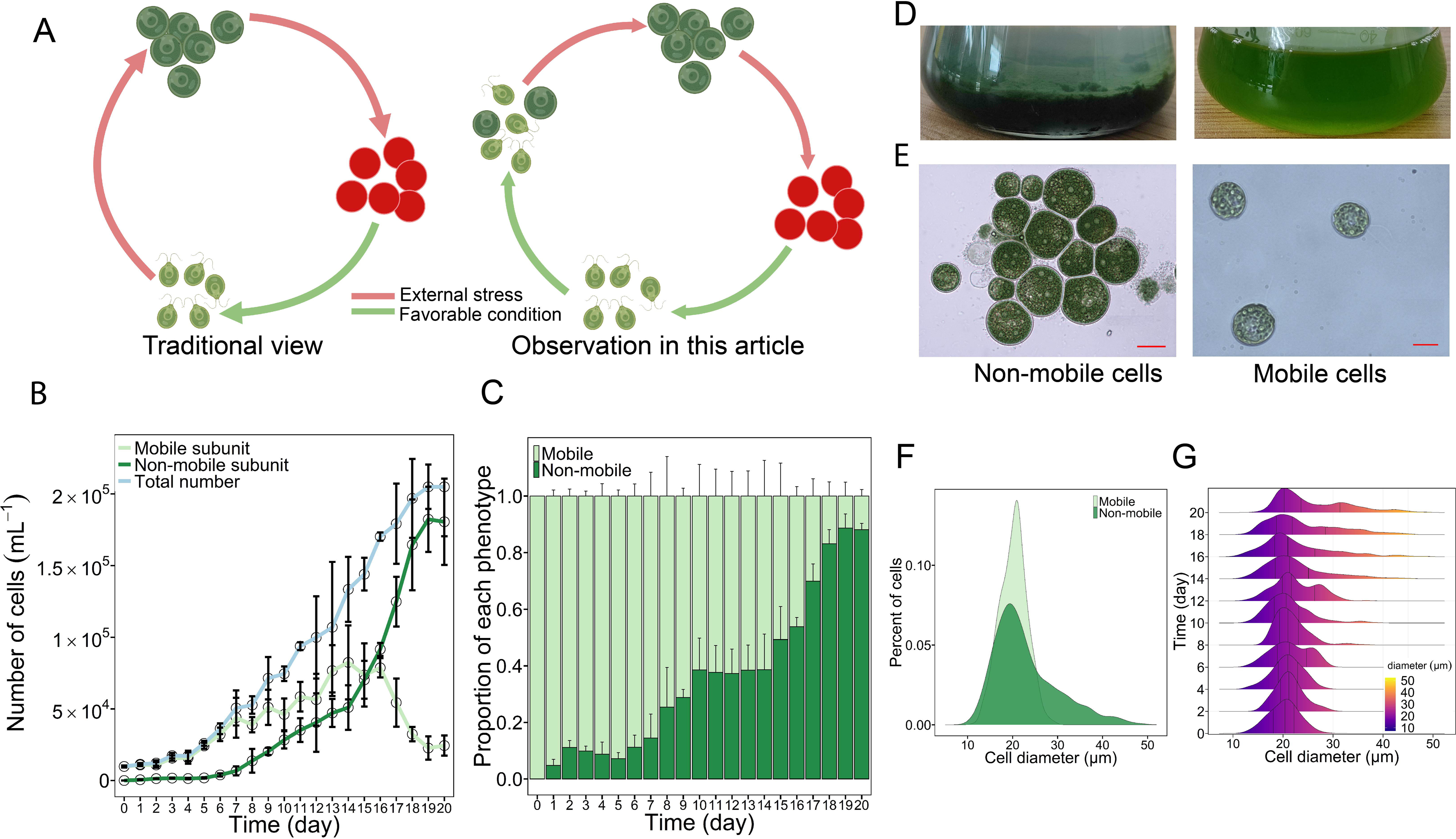
Phenotypic diversification of *H. pluvialis*. **A** Graphical summary of the traditionary view (left) and the main observation in this study (right). **B** Growth curve and phenotypic dynamics over the 20-day period. The blue line shows the total number of cells, the light green and dark green lines show the number of mobile cells and non- mobile cells, respectively. The error bars indicate the standard deviation of the triplicate values. **C** A bar chart showing the change in the ratio of mobile (light green) and non- mobile phenotype (dark green) over the period of 20 days. The error bars indicate the standard deviation of the triplicate values. **D** Images of enriched non-mobile cells (left) and a natural mobile population (right). **E** Microscopic images of non-mobile (left) and mobile cells (right) (Scale bar: 20 μm). **F** The Gaussian Kernel Density curve illustrating the diameter distribution of mobile (light green) and non-mobile cells (dark green). **G** The Gaussian Kernel Density curve showing the distribution of diameter on every other day of the growth experiment over the period of 20 days.

## Results

### Phenotypic diversification of *H. pluvialis*

First, a population starting with 100% mobile *H. pluvialis* cells was grown in fresh BBM medium without external stress. After 24 hours, the differentiation into a hetero- phenotypic population, composed of mobile and non-mobile cells, could be detected (Fig. 1B & Fig.1C). As the mobile cells converted into non-mobile cells, the ratio of mobile cells decreased, while the percentage of non-mobile cells increased over time. Specifically, the percentage of mobile cells decreased from 100% to 8.8%, while the non-mobile cells accounted for 91.2% of the total population at the end of the experiment (Day 20, Fig.1C). It is interesting to note the early occurrence of the non-mobile phenotype, which already accounted for almost 10% of the total population on day 3, when it is unlikely that there is any nutrient or space stress. This is also reflected when measuring the remaining essential nutrients of the medium, NO_3-_ and PO_43-_, which amounted to 96.7% and 83.7% of their original levels, respectively, on day 5 and to 47.3% and 35.6% of the original levels each on day 20 (Fig. S1). Since phenotypic diversification occurred shortly after inoculation, and cell density was relatively low, it can be assumed that no spatial stress was present either.

Mobile and non-mobile cells differ in both behaviour and morphology. Visible to the naked eye, non-mobile cells lost mobility and settled to the bottom of the Erlenmeyer flask with a layer of translucent medium on top (Fig. 1D, left), which is clearly different to a mobile population with a homogenous green color and where no precipitation is observed (Fig. 1D, right). In contrast to the free-living swimming lifestyle of the mobile cells (Fig. 1E, right), we also observed under the microscope that non-mobile cells lost two flagella and most cells aggregated into clumps (Fig. 1E, left).

Another important trait is cell size, changes in which will have profound effects on cell physiology. We first compared cell size between mobile and non-mobile cells and then recorded the dynamics of cell size along with the growth experiment. Our results showed an overlapping size distribution between the two phenotypes (Fig. 1F).

Specifically, mobile cells showed a less variable size distribution, ranging from 15 µm to 25 µm in diameter. The non-mobile population generally showed an increasing trend in cell size (15 µm to 50 µm), with 29.1% of cells having a diameter greater than 25 µm and some cells up to 50 µm in diameter.

The cell size dynamics of the growth experiment, with more and more mobile cells converting to non-mobile cells over time, showed a trend of gradual increase in overall population size, with about 25% of the population having a cell size diameter larger than 30 µm at day 20 (Fig. 1G).

### Tradeoff between growth and survival

To test whether the observed heterogeneity in *H. pluvialis* morphology and behaviour is bet hedging, we first investigated whether there is a tradeoff between growth, i.e., increase in cell numbers, and survival, i.e., stress resistance at the population level. Accordingly, we tested and compared cell numbers of four populations, i.e., mobile cell-only (Mobile), non-mobile cell-only (Non-mobile), two artificially created “bet-hedging” populations (20% mobile and 80% non-mobile cells, hereafter M2NM8; 80% mobile and 20% non-mobile cells, hereafter M8NM2), under standard culture conditions, i.e., supernatants from day 10 and under stress conditions (salinity stress, oxidative stress and drought stress) over time. Standard culture conditions leading to lower cell numbers and stress conditions leading to higher cell numbers of the non-mobile population would provide evidence that non-mobile cells trade growth for stress resistance and thus function as the persistent subunit of the population.

Under standard conditions, we observed a 7.9-fold higher cell count in the Mobile population than in the Non-mobile population in an 11-day experiment. The cell numbers of M8NM2 and M2NM8 were lower than those of the Mobile population, but 5.1-fold and 2-fold higher than those of the Non-mobile population, respectively (Fig. 2A). The variance in cell numbers between two artificial bet-hedging populations manifests the trend that the higher the proportion of mobile cells, the higher the cell numbers.

**Figure. 2.**
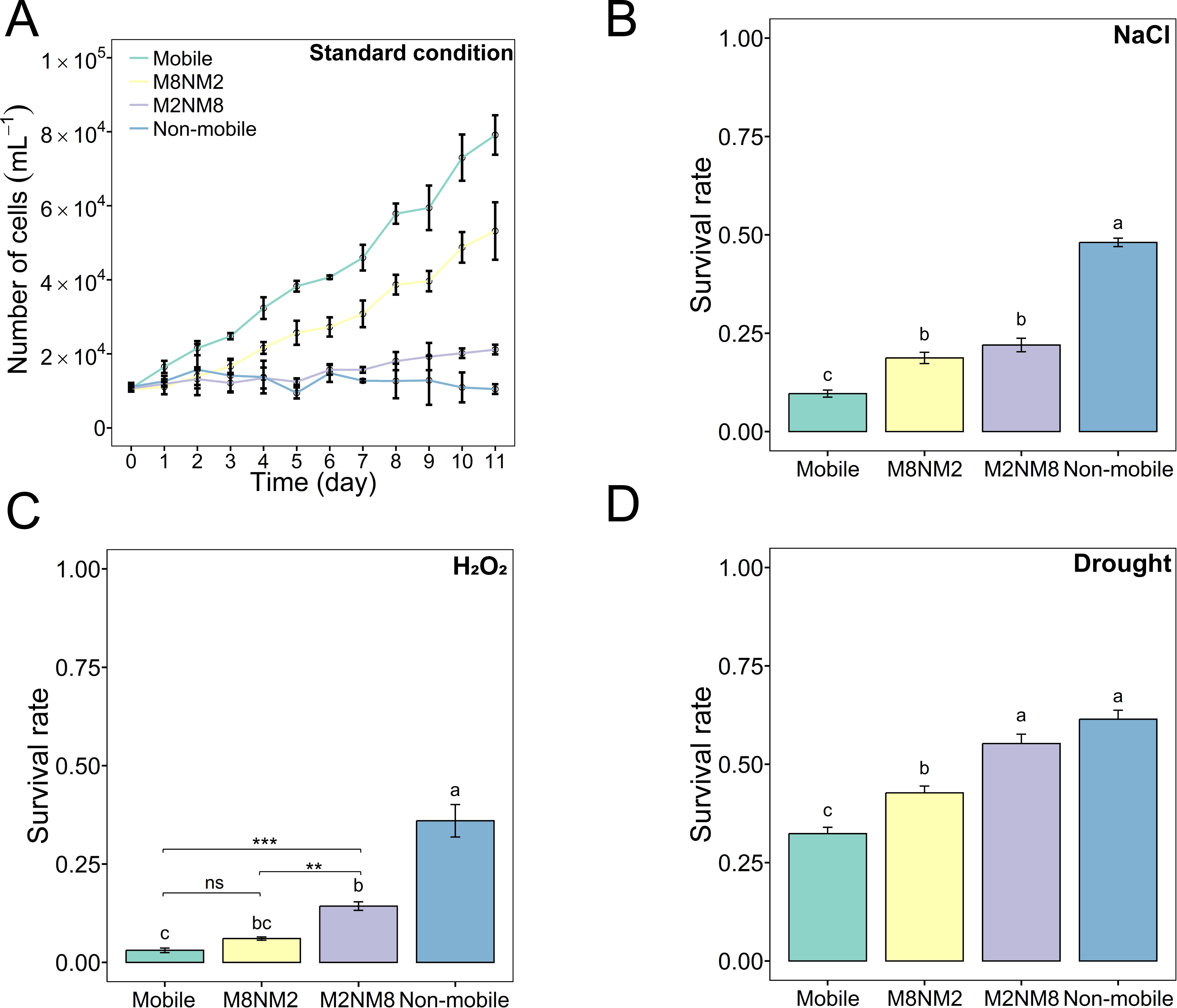
Growth and survival of each population in response to standard and adverse conditions. Growth and survival of four populations of different phenotypic composition were evaluated, i.e. Mobile, M8NM2, M2NM8, Non-mobile. **A** Growth of four populations under standard conditions over the period of 11 days. **B** Survival rate of four populations under salinity stress (300 mM NaCl). **C** Survival rate of four populations under oxidative stress (1 mM H_2_O_2_). **D** Survival rate of four populations under drought stress (immediate desiccation). All three survival rate assays under stress conditions lasted for 10 days. Error bars indicate standard deviation of triplicate values. Different letters on each graph represent statistical significance (*p* < 0.05), calculated by one-way ANOVA with Tukey’s HSD post-hoc analysis across all four populations. Line segments and asterisks in **C** indicate statistical significance calculated by one-way ANOVA without the Non-mobile treatment. Mobile (mobile cells only, green), M8NM2 (80% mobile and 20% non-mobile cells, yellow), M2NM8 (20% mobile and 80% non-mobile cells, purple), and Non-mobile (non-mobile cell only, blue). Significance: ns (no significance), *(*p* < 0.05), **(*p* < 0.01), ***(*p* < 0.001), ****(*p* < 0.0001).

Although the four populations behaved differently under different environmental stresses, there was a general tendency for the Non-mobile population to have the highest survival, the Mobile population to have the lowest survival, and the survival rate of the two artificial bet-hedging populations was in between. When exposed to high salinity (300 mM NaCl) for ten days, the Non-mobile population showed the highest survival rate (48.1%), followed by the two hetero-phenotypic populations, 22% for M2NM8 and 18.7% for M8NM2, respectively. The Non-mobile population had an 8.6-fold fitness advantage over the Mobile population, which had the lowest survival rate of 9.7% (ANOVA, F_3,8_ = 156.2, *p* < 0.001, Fig. 2B). Hydrogen peroxide (H_2_O_2_) treatment was performed to represent oxidative stress, and all populations were more sensitive to oxidative stress than to salinity stress. After H_2_O_2_ exposure (1 mM), survival was highest in Non-mobile, at 36%. The M2NM8 population had the second highest survival rate (14.3%). In the M8NM2 and Mobile populations, almost all cells perished and the survival rate was 6% and 2%, respectively (Fig. 2C). Drought stress was better survived by all populations. Although significantly lower than the survival rate of the other three populations, 42.7% of the Mobile population survived. The Non-mobile population still performed best, with a survival rate of 61.4%, followed by 55.3% for M2NM8 and 42.7% for M8NM2 (Fig. 2D).

In summary, these assays provide direct evidence that: (1) the non-mobile phenotype traded growth for stress resistance; (2) the two hetero-phenotypic populations had lower variance in the two fitness-related traits under different conditions, always in the middle of two mono-phenotypic populations. Taken together, this evidence supports our hypothesis of bet hedging.

### Distinct physiological profiles of the two phenotypes

To explore the underlying mechanisms of the higher stress resistance of the non-mobile phenotype, we examined critical physiological parameters in both phenotypes, including photosynthetic activity, cellular dry weight, pigment, lipid and starch content. Photosynthesis was used as an indicator of stress response, as it is widely accepted that photosynthetic organisms reduce the energy budget in growth-promoting photosynthesis in exchange for investing more energy in activities associated with stress resistance^30,31^. The chlorophyll fluorescence parameter F_v_/F_m_ was tested to determine the maximum quantum efficiency of photosystem II (PSII), an indicator of photosynthetic activity. The result showed that non-mobile cells had a significantly lower F_v_/F_m_ value than mobile cells (*t* test, *p* < 0.05, Fig. 3A), indicating reduced photosynthetic activity. In addition, the pigments involved in photosynthesis, chlorophyll *a* and *b*, showed a significantly decreased pattern, accounting for 63.5% and 61.5% of the value in mobile cells, respectively. Another pigment, carotenoid, was also quantified, and showed an increasing pattern, about twice that of mobile cells (Fig. 3B).

**Figure. 3.**
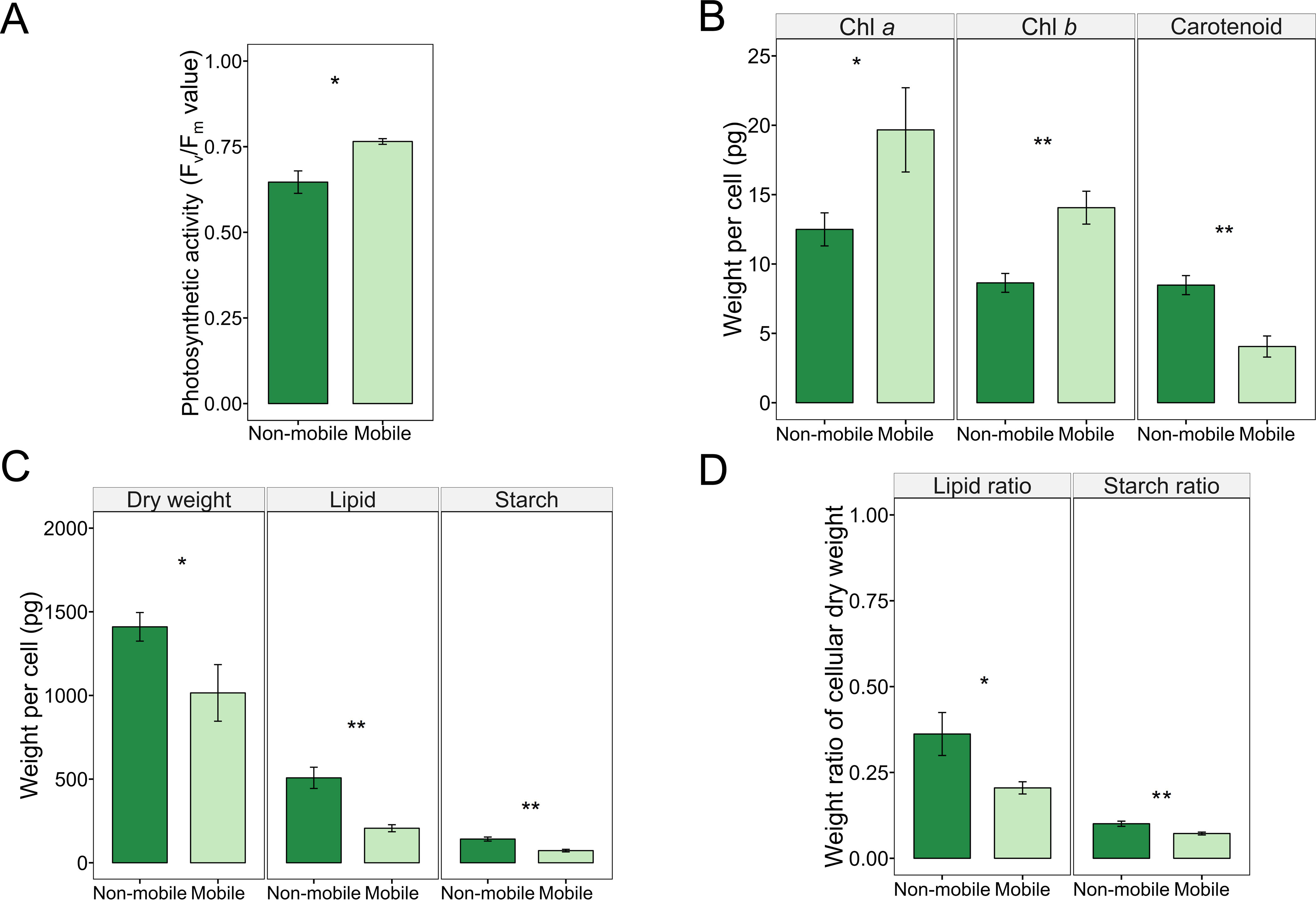
Physiological profiles of mobile and non-mobile phenotypes. **A** Chlorophyll fluorescence parameter F_v_/F_m_. **B** Cellular chlorophyll *a*, chlorophyll *b* and carotenoid content. **C** Cell dry weight, lipid and starch content. **D** The ratio of cellular lipid and starch to dry weight. Error bars denote the standard deviation of triplicate values. Asterixes represent statistical significance calculated by Welch’s *t* test. Mobile cells (light green), non-mobile cells (dark green). Significance: *(*p* < 0.05), **(*p* < 0.01), ***(*p* < 0.001), ****(*p* < 0.0001).

Lipids and starch are considered critical energy storage substances that play a crucial role in response to adverse environmental conditions^32–34^. Higher cellular levels of these substances in non-mobile cells will help us to understand how they achieve higher stress resistance. Cellular lipids, starch, as well as dry weight of non-mobile cells increased significantly by 1.39-fold (*t* test, *p* < 0.05), 2.46-fold (*t* test, *p* < 0.01) and 1.95 fold (*t* test, *p* < 0.01) (Fig. 3C), respectively, compared to mobile cells, indicating an accumulation of these metabolites. Furthermore, an increase in cellular dry weight indicates a shift in energy budget from cell growth to storage. In addition, the ratio of lipids (to dry weight) increased significantly from 20.5% to 36.2% (*t* test, *p* < 0.05), while the ratio of starch to dry weight rose from 7.2% to 10% (*t* test, *p* < 0.01) (Fig. 2D), suggesting that lipids are the primary energy storage substance and starch may have an anaplerotic function.

### Distinct transcriptomic profiles between phenotypes

To understand the underlying molecular background and to capture the transcriptomic characteristics of each phenotype, the respective cells were sequenced to examine the key differentially expressed genes (DEGs). In general, 6128 genes of the non-mobile cells showed statistically significant changes in mRNA level compared to the mobile cells (adjusted *P* value < 0.05), with the transcript level of 2953 genes increased and 3185 genes showing a decreased expression pattern (Fig. 4A). According to the results of the KEGG enrichment analysis, all DEGs were classified into several functional categories, showing striking metabolic differences, such as in photosynthesis, carotenoid biosynthesis, pyruvate metabolism, and fatty acid (a major component of lipids in microalga) and starch metabolism (Fig. 4B, top 20 pathways). Particular attention was paid to the expression level of specific genes of four major metabolic pathways, i.e. photosynthesis, carotenoid, lipid and starch metabolism, which are thought to be associated with the cellular stress response. The expression patterns of key genes involved in these four metabolic pathways could clarify at the molecular level how the non-mobile phenotype proves to be more stress-resistant.

**Figure. 4.**
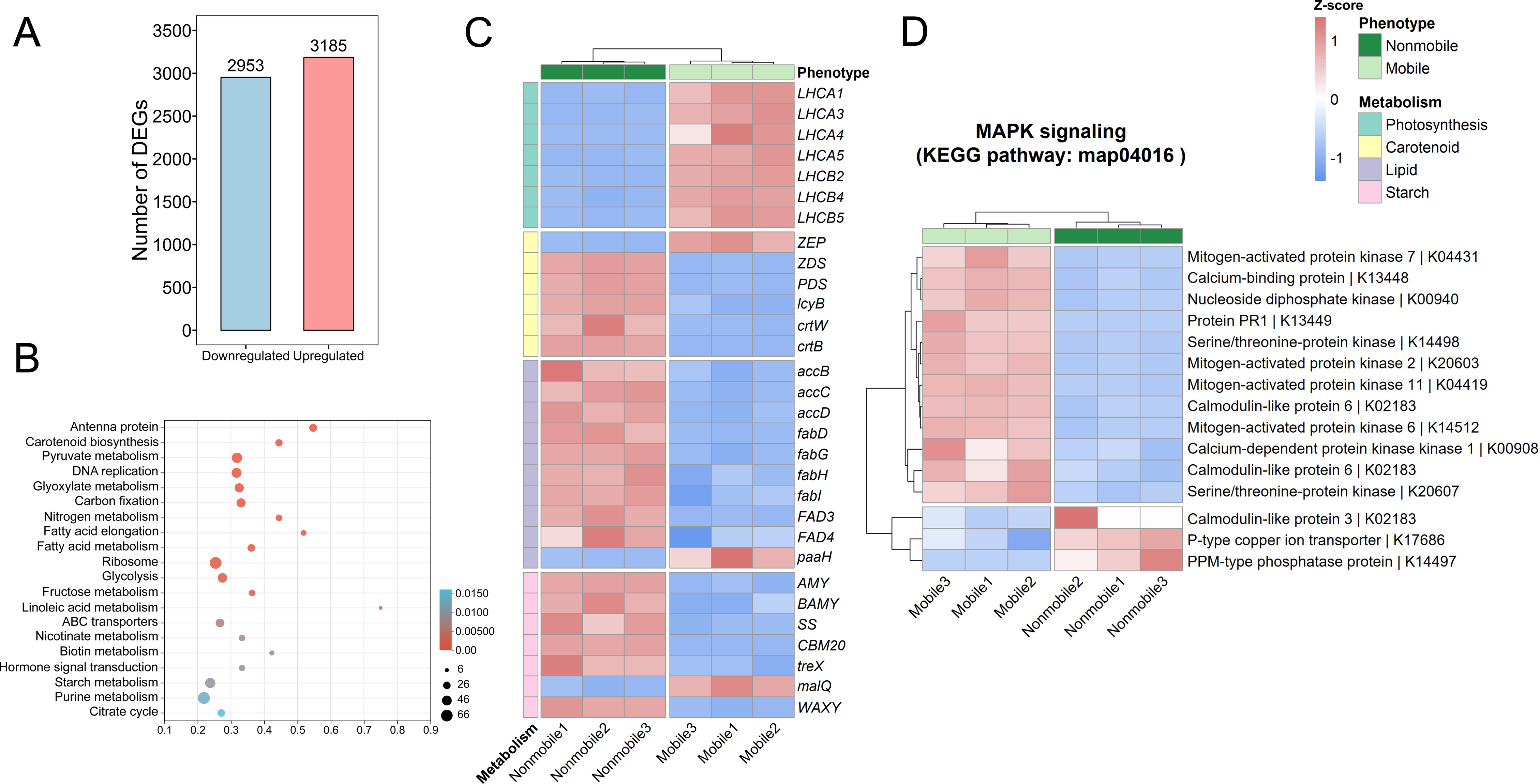
Transcriptome and core functional enrichment analysis of differentially expressed genes. **A** Bar chart of differentially expressed genes. The numbers on the top of the bar indicate the number of genes that were significantly up- or down-regulated. **B** Dot plot analysis of enriched KEGG Pathways with differentially expressed genes with *p* value < 0.05. The size of the dot indicates the number of genes for each pathway, and the color bar indicates the *p* value. **C** Heatmap analysis showing DEGs enriched for four KEGG pathways, including photosynthesis, carotenoid, lipid, and starch metabolism. Phenotype treatments are non-mobile (dark green) and mobile (light green). The metabolic treatments are Photosynthesis (blue), Carotenoid (yellow), Lipid (purple), and Starch (pink). The heatmap color gradient shows low gene expression (blue) and high gene expression (red). **D** Functional enrichment analysis of the MAPK signaling pathway. Heatmap analysis of DESeq2-normalized gene expression scaled as the number of standard deviations from the row means for all genes with KO terms under the KEGG MAPK signaling pathway (map04016). KEGG annotations were assigned from the genome annotation. Column dendrograms show similarity based on Euclidean distance and hierarchical clustering. Gene clusters were determined by k-means clustering with Euclidean distance.

In detail, in photosynthesis, genes involved in the synthesis of antenna proteins (a set of important proteins for the assembly of the light-harvesting complex), such as LHCA 1, 3, 4, 5 and LHCB 2, 4, 5^35,36^, showed a decreased pattern in non-mobile cells, indicating reduced photosynthetic activity. The decreased transcription levels of the antenna proteins are consistent with the F_v_/F_m_ assay, where the non-mobile cells had significantly lowered F_v_/F_m_ levels. As for carotenoid metabolism, critical genes involved in carotenoid biosynthesis were upregulated, while genes associated with carotenoid degradation showed a decreasing pattern. For example, the core carotenoid biosynthesis genes *crtB, crtW, lcyB, PDS* and *ZDS*^37–39^ showed significantly higher expression levels. Instead, the expression of ZEP (zeaxanthin epoxidase), an important carotenoid regulation gene^39,40^ that contributes to the maintenance of normal carotenoid levels, decreased. A similar transcriptomic pattern was observed in lipid metabolism. Two prominent lipid biosynthesis gene families, *acc* (*accB, accC, accD*) and *fad* (*fadD, fadG, fadH, fadL, fad3, fad4*)^41^, were significantly upregulated, while the lipid degradation gene *paaH*^42^ was downregulated. The transcription level of related genes suggests that non-mobile cells accumulate carotenoids and lipids. The starch synthase *SS*^43^ and the gene *WAXY*^44,45^, associated with starch granule biosynthesis, were markedly upregulated, suggesting accumulation activity. Surprisingly, we also found that several genes involved in starch degradation (*AMY, BAMY, CBM20, treX, malQ*)^43,46–49^ significantly increased their transcript levels.

Furthermore, the transcriptomic levels of mitogen-activated protein kinase (MAPK) genes are evaluated to explore whether non-mobile cells are induced by external stressors. MAPK cascades are evolutionarily conserved signaling modules that serve to convert environmental stimuli into intracellular responses^50^. Various stressors such as temperature, drought, salinity, UV irradiation and reactive oxygen species can activate MAPK signalling pathways^51^. Cellular behaviors regulated by MAPKs are thought to play an essential role in defence responses^50–52^. The expression pattern of the annotated MAPK genes (12 out of 15 annotated genes show a relatively down- regulated pattern of the non-mobile phenotype, Fig. 4D) illustrates that the transformation of the non-mobile cells was probably not caused by stress to which the mobile cells were exposed, at least not on the day of sampling (day 10). Given the transformation into the non-mobile state takes time, those transformed ancestor mobile cells at earlier time points should not be stressed either. This result suggests that the formation of non-mobile cells and accompanying physiological modifications may occur stochastically (stochastic gene expression), another feature of the bet hedging theory^53^.

### Ability of both phenotypes to re-diversify

To distinguish between “bet hedging”, “evolutionary radiation”^54^, “genetic mutation”^55,56^ and “cellular age”^57,58^, all of which can lead to phenotypic diversification, we determined whether mobile and non-mobile cells could re-diversify when re-cultured for another round of growth. The results showed that the mobile population diversified back into a hetero-phenotypic population. In addition, the non-mobile cells reverted to a mobile status and first regained fertility, then diversified back to the non-mobile phenotype over time (Fig. 5A, Fig. 5B). The phenotype of the founding cells also had no effect on this diversification: both populations initiated with mobile and those initiated with the non-mobile phenotype formed diversified phenotypes, with non-mobile cells constituted averaging 20.6% and 18.9% of the total, respectively (Fig. 5B). These data suggest that the emergence and re-emergence of phenotypic diversification are not due to evolution, mutation or ageing but to bet hedging.

**Figure. 5.**
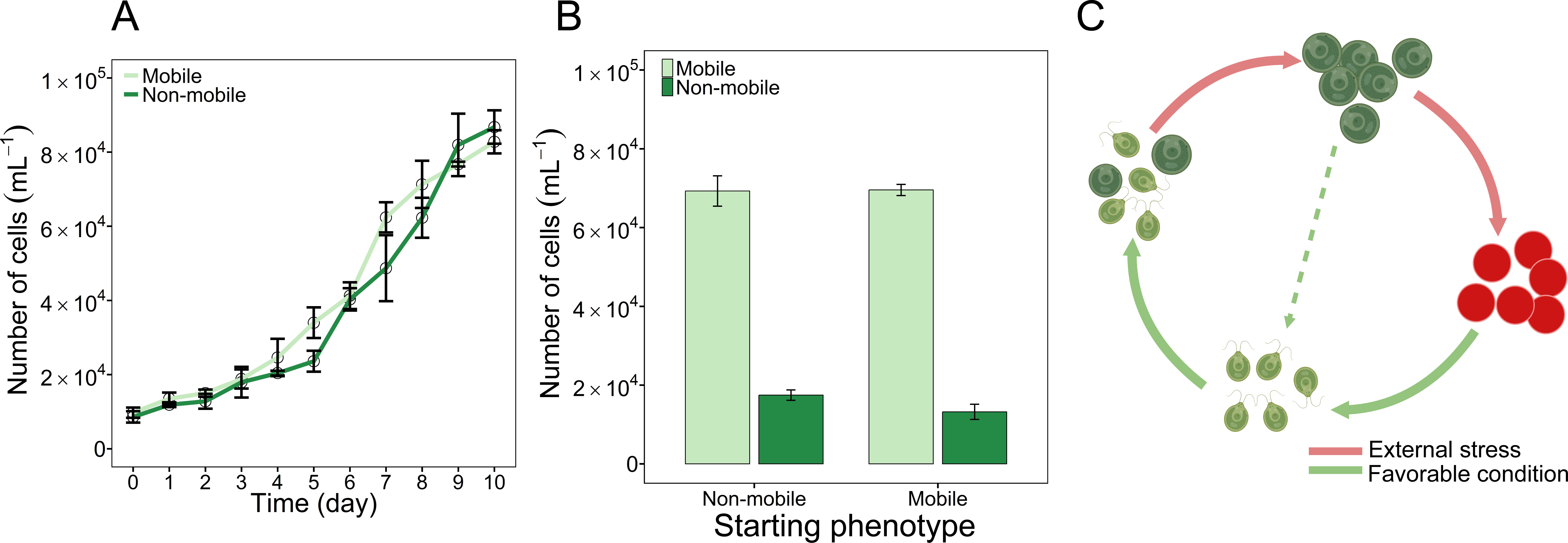
Rediversification of the mobile and non-mobile phenotype. **A** Growth of the population consisting of only mobile or non-mobile cells in fresh BBM over the period of 10 days. **B** Bar chart of population composition on day 10. Error bars indicate standard deviation of triplicate values. Mobile cell population only (light green), non-mobile cell population only (dark green). **C** Graphical representation of the reversibility of the non-mobile phenotype to a mobile state (dashed line).

## Discussion

Unlike other adaptive strategies, such as adaptive tracking and phenotypic plasticity^59,60^, bet hedging offers an alternative way to adapt to randomly fluctuating environments. For a genetically clonal population, it might be advantageous to hedge the bet, population fitness, by differentiating into different phenotypes. The assumed advantage of possessing different phenotypes with different survival chances and reproductive success relative to each other is that they are best suited to different conditions. Consequently, a subpopulation already exhibits the appropriate phenotype for the new environment before the external environment changes, ensuring that at least a subpopulation can survive and reproduce successfully. Currently, bet hedging is frequently reported to be a common adaptative strategy in various organisms. However, to our knowledge, it has never been reported and experimentally verified in microalga, a key player in aquatic ecosystems.

Due to the property of *H. pluvialis* to produce astaxanthin, this Chlorophyta microalga has attracted much attention. Interestingly, it has been repeatedly reported that the green stage of this species consists of two phenotypically distinct types at the population level: mobile and non-mobile^24–26^. Such phenotypic heterogeneity has been described for a range of other systems^61–64^ and is mainly thought to result from adaptation to microscale environmental variation^53,65–68^.

In the case of *H. pluvialis*, it has been assumed that non-mobile cells arose from mobile cells due to external stresses^26,69^. This might be important for older cultures due to increasing nutrient and space stress and can potentially be observed in our study as an increasing proportion of non-mobile cells over time (Fig. 1B & Fig. 1C). However, this does not explain the early appearance of non-mobile cells at the beginning of the culture period, suggesting that this phenotypic diversification is not triggered by stress at least at these early time points. This irrelevance is also supported by the relatively downregulated MAPK pathway genes (stress-dependent signaling pathway) of the non-mobile cells (Fig. 4D), suggesting that the non-mobile cells do not suffer from stress and may not have arisen from a response to stress either.

Another explanation for the observed phenotypic heterogeneity of *H. pluvialis* is “bet hedging”, in which the mobile cells represent the fast-growing subunit and non-mobile cells represent the stress-resistant subunit of the population. At the expense of reduced population growth caused by the transformation of some cells to a slow/non-growing, non-mobile phenotype (given the high proportion of small (less than 20 µm in diameter), non-mobile cells at day 20 (Fig. 1F), further evidence is needed to clarify whether non-mobile cells can reproduce or not), bet-hedging populations maintain a balance between population growth and stress resistance, ensuring both population reproduction and population survival that is not zero.

Apart from the notable difference in mobility, the two phenotypes also differ in other traits such as cell size and grouping behavior. Non-mobile cells generally have a larger cell diameter, and in contrast to free-living mobile cells, most of them aggregated into clumps. Such aggregation behavior could serve to establish a stable microenvironment or defend against size-dependent predators. In addition, although not investigated in our study, non-mobile cells have been reported to form a new cell wall^70^ that, together with the original wall, acts as a physical barrier, protecting some of the cells from external harmful chemical exposure or physical damage.

We further evaluated population growth and survival as indicators of fitness in four artificially created populations that differed in their ratio of mobile to non-mobile types under standard and stress conditions. The results were in line with our expectations, as: (1) the non-mobile phenotype traded growth for stress resistance, and (2) both “bet-hedging” populations showed lower variance in the fitness-related parameters across environments (Fig. 2). Furthermore, the ability to re-diversify of each phenotype suggests that the phenotypic heterogeneity in this study is due to bet hedging rather than evolutionary radiation, mutation or ageing; a similar phenomenon was observed in another bet-hedging bacterium *Sinorhizobium meliloti*^15^. Overall, our results provide clear evidence that the phenotypic heterogeneity of *H. pluvialis* serves as a bet-hedging strategy.

It is of general interest to understand how the non-mobile subpopulation becomes more stress-resistant. In the study of microalga for stress resistance, it is well documented that persistent cells typically reduce photosynthetic output and other non-essential metabolic activities, and instead enhance metabolic processes associated with persistence. For example, the accumulation of antioxidative substances as well as energy storage materials, including lipids and starch, is thought to be critical for resistance to environmental stress^39,71–74^. Accordingly, we examined the physiological and transcriptomic profiles of the non-mobile phenotype, with a focus on these metabolites, to uncover the underlying mechanisms of stress resistance. In general, the integrated physiological and transcriptomic data show that non-mobile cells have shifted from growth-promoting photosynthesis to resistance-promoting activities, including the accumulation of antioxidant and energy-storing substances.

In detail, our results show that non-mobile cells significantly decreased photosynthetic activity, based on the F_v_/F_m_ assay (Fig. 3A), which might result from the decreased amount of chlorophyll *a* and *b* (Fig. 3B) and the decreased expression of light-harvesting complex proteins (Fig. 4C). Instead, cellular carotenoids (antioxidative substances)^75^, lipids and starch (energy storage compounds)^32^, all three increased significantly, as expected (Fig. 3B & Fig. 3C). This is confirmed by the transcriptomic analyses and suggests a tradeoff between growth and stress resistance. Counterintuitively, five genes related to starch degradation (*AMY, BAMY, CBM20, treX, malQ*) also showed higher expression in non-mobile cells (Fig. 4C), making the results intriguing to interpret. We propose that the accumulation of starch could serve as a rapid response to stress and that starch acts as an intermediate energy storage compound, which could later be used to supply energy to other subsequent processes, such as lipid biosynthesis, which is considered long-term energy storage. This “confusing” gene expression pattern could explain why lipids accounted for 36.2%, but starch only for 10% of the cell dry weight (Fig. 3D). Together with the near arrest of growth, increased cell size and aggregation behavior (Fig. 1E & Fig. 1F), all traits provide evidence to explain how non-mobile cells are more stress resistant than their mobile counterparts.

More interestingly, in contrast to other reports of microbial bet hedging, where only one stress-resistant phenotype exists, in *H. pluvialis*, the final resistant form is the red aplanospore^23,26^. Non-mobile green cells continue to develop into aplanospores if the environmental conditions deteriorate further. Red aplanospores are thought to be able to survive extremely harsh environmental conditions, although here we show that the non-mobile cells are already resistant to stress. The existence of two different resistance phenotypes would make sense from an evolutionary point of view, as the non-mobile green cells represent a fast response to the environment and can quickly revert to a fast-growing state, i.e., the mobile phenotype, once conditions improve (Fig. 5). If environmental conditions continue to deteriorate, the non-mobile green cells transform into aplanospores, the ultimate resistant form that can survive harsher conditions for longer. The presence of two progressive persistence phenotypes could serve as an essential and effective adaptation strategy for *H. pluvialis* in nature, which would have been selected by the species’ complex natural habitats. Indeed, unlike microalgal species living in open waters, the natural habitats of *H. pluvialis* are often small, temporary natural or artificial aquatic habitats^76–80^, whose conditions can easily change and dramatically alter, posing extreme challenges for the organisms living in them. Under such conditions, the survival and reproduction of the *H. pluvialis* population and subsequent competition for nutrients and space would particularly favor bet hedging. By forming mobile and non-mobile subgroups, the *H. pluvialis* population could take advantage of changing environmental conditions that can be devastating for other microalgal species, while increasing the chances of survival until the environment is favorable again.Similar to the difficulties in identifying bet-hedging behaviors in bacteria, determining microalgal bet hedging is relatively challenging, as identifying phenotypic diversification or metabolic variation and then isolating the corresponding phenotypes of a population might be difficult. This is likely why, although this has been occasionally suggested^81^, to date, there are no experimental studies that provide solid experimental evidence for bet hedging. In nature, however, bet hedging of microalga should occur more widely and frequently than expected. For example, some blooming microalgal species that form resting cysts as part of their life histories, can not only survive global environmental changes, but also thrive even in the face of human interventions, such as physical, chemical, and biological anti-blooming attempts. To some extent, this can be attributed to the unreported “bet hedging” of these species, i.e., the simultaneous co-existence of vegetative cells and cysts that allow them to survive anthropogenic disturbances and re-prosper again.

In summary, here, we provide evidence that the observed phenotypic heterogeneity in *H. pluvialis* is a bet-hedging strategy, in which parts of the mobile population transform spontaneously and reversibly into a distinct, non-mobile state that exhibits almost no growth but significantly higher resistance to stressors. Measurements of fitness proxies, i.e., population growth and survival rates, show that hetero-phenotypic populations have reduced variance in these parameters under different conditions, which aligns with the basic concepts of bet hedging. Further analyses indicate that the ability of the non-mobile phenotype to withstand stress is due to metabolic shifts from growth-promoting photosynthesis to stress-resistant activities such as cell enlargement and aggregation, as well as the accumulation of antioxidative substances (carotenoids) and energy storage compounds (lipid and starch). However, without a record of the long-term frequency and magnitude of stressful conditions faced by *H. pluvialis* in the field, we cannot conclusively determine the evolutionary and molecular origins of the phenotypic heterogeneity, this limitation is nearly ubiquitous in studies of bet hedging. Overall, this study provides the first experimental account of “bet-hedging” behaviour in the phyla of microalga, providing fundamental insights into how these organisms can cope with unpredictable, fluctuating environments and offering a different perspective for future species or ecosystem rescue or management, and prompting us to rethink the mass-cultivation strategy of this commercially valuable species.

## Methods

### Strain and culture condition

*Haematococcus pluvialis* (FACHB-712, Freshwater Algae Culture Collection at the Institute of Hydrobiology, Wuhan, China), also known as *Haematococcus lacustris,* was maintained photoautotrophically in freshwater BBM medium as previously described^82^. Specifically, the stock culture was maintained in 500 mL of freshwater medium BBM in Erlenmeyer flasks at 25±1°C under 12h/12h light-dark cycles (20 μmol photons m^−2^ s^−1^). The cells were grown without forced aeration, the only CO_2_ supply being diffusion from the atmosphere into the flasks.

### Growth experiment, phenotype and cell size dynamics

Prior to the growth experiment, the stock culture was first transferred to fresh BBM, and the mobility of the population was checked daily under an inverted microscope (Primo Vert, Carl Zeiss, Germany). From experience, a high proportion of cells (> 95%) show the ability to swim in the first few days when transferred to a new medium, even if it was initiated with a hetero-phenotypic population composed of mobile and non-mobile mother cells. Because non-mobile cells lose their swimming ability and will sediment to the culture flask bottom, by gently pipetting the upper-level culture after standing still for one hour, we can enrich the mobile cells. In this way, a 100% mobile population was achieved, which was used in the subsequent growth experiment.

In the growth experiment, an initial density of about 1*10^4^ cells/mL of cells was inoculated into 200 mL of fresh medium in an Erlenmeyer flask at 25±1°C under 12h/12h light-dark cycles (20 µmol photons m^−2^ s^−1^) for 20 days in triplicate. Cell density and population composition based on mobility were recorded daily. Specifically, 100 µL culture was sampled after careful mixing and counted with a counting chamber under an inverted microscope. Mobile cells and non-mobile cells were distinguished from each other and recorded on the basis of observing the mobility of an individual after 5 minutes of accommodation in the counting chamber to exclude the effects of physical disturbance during sampling.

Meanwhile, another 100 µL culture was used to measure cell size. A thin layer of low melting point agarose gel (Thermo Fisher Scientific, USA) was prepared to stabilize the cells. Images were then captured and processed using AxioVison SE64 Rel.4.9 software, where cell diameters were determined using the length measurement function. For each replicate at each time point, cell diameters of approximately 300 to 500 cells were measured.

### Separation of subpopulations and determination of cell size

Due to the precipitating and clustering properties of the non-mobile phenotype, the two cell types are relatively easy to isolate and enrich. For the isolation of the mobile cells, 10-day-old populations of the growth experiment were kept still for one hour to allow the non-mobile cells to settle to the bottom, and the upper green part of the culture, containing the mobile cells was pipetted out and enriched into new flasks. For the non-mobile cell isolation, the remaining mobile cells were resuspended in 20 mL of added double distilled water (ddH_2_O, ipureplus, neoLab Migge, Germany), after which the liquid was pipetted out and discarded. Another 20 mL of ddH_2_O was used to wash off the remaining non-mobile cells by pipetting. The separate mobile and non-mobile cultures were checked under the microscope to ensure that the two phenotypes did not coexist. To determine the cell size of each phenotype, the cell diameter of the two separate cultures was examined under the microscope as described above.

### Nutrient dynamics over time

Residual inorganic nutrients (nitrate and phosphate) of the culture medium were measured every five days using the Auto Discrete Analyzer (Cleverchem Anna, Dechem-Tech, Germany). In particular, nitrate and phosphate concentrations were measured using the cadmium-zinc reduction method and molybdenum-antimony spectrophotometry, respectively^83^.

### Microscopy

Microscopic images of mobile and non-mobile cells were taken using an inverted fluorescence microscope (Axio observer Z1, Carl Zeiss, Germany).

### Quantification of cell dry weight, chlorophyll, carotenoids, starch and lipids

For the determination of cell dry weight, 10 mL of culture was collected in a pre-dried centrifuge tube and centrifuged at 6,000 rpm for 5 min (Eppendorf, Germany). The cell pellets were washed three times with deionized water, dried for three hours at 105°C in an oven (Yiheng, Shanghai, China) and then cooled to room temperature before weighing.

Chlorophyll *a*, chlorophyll *b* and carotenoid contents were measured according to a previously published method^84–86^. In brief, 10 mL of the culture of each phenotype was centrifuged at 14,000 rpm for 10 min at 4°C. The supernatant was discarded, and the remaining cells were resuspended in 2 mL of 90% methanol (Aladdin, Shanghai, China). The suspension was incubated in a water bath at 70°C for 10 mins and centrifuged again at 14,000 rpm. The supernatant was used to measure absorbance measurement with an ultraviolet-visible (UV–Vis) spectrophotometer (UV-1780, SHIMADZU, Japan) at wavelengths of 470, 652 and 665 nm. The pigment content was determined using the following formulas: Chlorophyll *a* (mg L^-1^) = 16.82 A_665_ - 9.28 A_652_; Chlorophyll *b* (mg L^-1^) = 36.92 A_652_ - 16.54 A_665_; Carotenoid (mg L^-1^) = (1000 A_470_ - 1.19 C_a_ - 95.15 C_b_) / 225; where A_470_, A_652_ and A_652_ are the absorbances at wavelengths of 470, 652 and 665 nm, respectively.

Cellular starch was quantified according to a published method with some modifications^87^. 3 mL of each sample was ground with a grinding rod. The pigments were extracted three times with 4 mL of 80% ethanol for 15min at 70°C. For complete hydrolysis of the starch, 0.6 mL of 52% perchloric acid was added to the precipitate, stirred for 30 min at 25°C and centrifuged. This procedure was repeated three times. Then 0.2 mL of the extract was cooled to 0°C; 1 mL of anthrone solution [0.1 g of anthrone in 50 mL of 82% (v/v) H_2_SO_4_] was added and stirred. The mixture was kept in a water bath at 100°C for 8 min and then cooled to 20°C. The absorbance was measured at 625 nm. Calibration was performed simultaneously with glucose as the standard.

The intracellular lipid content was determined with a neutral lipid-specific dye, Nile Red (9-diethylamino-5H-benzo(a)phenoxazine-5-one)^88^. Specifically, lipid content was determined by breaking cells with a grinding rod followed by extraction with chloroform/methanol (2:1, v/v) and total lipid content was measured gravimetrically. As for the samples, 40 µL of Nile Red solution in acetone (250 mg/L) was added to 1 ml of algae suspension (20% DMSO). The mixture was vigorously agitated with a vortex mixer. Fluorimetric analysis was performed 10 min after staining using an Enzyme labelling apparatus (Tecan, Swiss) with a 490 nm narrowband excitation filter and a 585 nm narrow band emission filter.

### Transcriptomics

100 mL of each phenotype was harvested as described above and centrifuged for 10 min at 4000 rpm at 4°C to remove the supernatant. 100 mL of ddH_2_O was added to resuspend the cell pellets and centrifuged again; the supernatants were discarded. This procedure was performed twice. The cell pellets were transferred to RNase-free cryogenic vials and frozen in liquid nitrogen for 30 min until extraction. In total, both cell types were analyzed in triplicate.

Total RNA was extracted with Plant RNA Purification Reagent for plant tissue according to the manufacturer’s instructions (Invitrogen, USA), and genomic DNA was removed using DNase I (TaKara, Dalian). Total RNA samples were sent to Shanghai Majorbio Bio-pharm Technology Co., Ltd. (Shanghai, China). RNA quality was determined using a 2100 Bioanalyzer (Agilent) and quantified using the ND-2000 (NanoDrop Technologies). Only high-quality RNA samples (OD_260/280_ = 1.8∼2.2; OD_260/230_ ≥ 2.0; RIN ≥ 6.5; 28S:18S ≥ 1.0; and > 1μg) were used for sequencing library construction.

The RNA-seq transcriptome library was prepared from 1 μg total RNA using the TruSeq^TM^ RNA sample preparation kit from Illumina (San Diego, CA, USA) using. Data were analyzed using the Majorbio Cloud online platform (www.majorbio.com). Raw paired-end reads were trimmed and quality controlled by SeqPrep (https://github.com/jstjohn/SeqPrep) and Sickle (https://github.com/najoshi/sickle) using the default parameters. The clean reads were then separately aligned to the reference genome with orientation mode using HISAT2 software (http://ccb.jhu.edu/software/hisat2/index.shtml). The mapped reads from each sample were assembled using StringTie(https://ccb.jhu.edu/software/stringtie/index.shtml?%20t=example) in a reference-based approach.

To identify the differentially expressed genes (DEGs) between two different treatments, the expression level of each transcript was calculated using the transcripts per million reads method (TPM). RSEM (http://deweylab.biostat.wisc.edu/rsem/) was used to quantify gene abundances. Essentially, differential expression analysis was performed using DESeq2 with a Q value ≤ 0.05, and DEGs with |log2FC| > 1 and a Q value ≤ 0.05 were considered significant DEGs. In addition, functional enrichment analyses, including Kyoto Encyclopedia of Genes and Genomes (KEGG) analyses, were performed to determine which DEGs were significantly enriched in KO terms and metabolic pathways at Bonferroni-corrected P-value ≤ 0.05 compared to the whole transcriptome background. Analyses of functional GO enrichment and KEGG pathways were performed by Goatools (https://github.com/tanghaibao/Goatools) and KOBAS (http://kobas.cbi.pku.edu.cn/home.do), respectively.

### Photosynthetic activity

Quantification of photosynthetic performance based on chlorophyll fluorescence was analyzed using the FL3500 fluorometer (PSI, Czech Republic). 2 mL of separated mobile and non-mobile cultures were placed in darkness for 20 mins, and the maximum quantum efficiency of PSII photochemistry (F_v_/F_m_) was measured immediately after dark adaptation using a PPFD of 3000 μmol m^−2^ s^−1^ as a saturating flash for 1 s duration.

### Growth and survival assays under various conditions

Four populations were included in fitness assays, i.e. mobile cell-only population (Mobile), non-mobile cell-only population (Non-mobile), as well as two artificially prepared bet-hedging populations (20% mobile and 80% non-mobile cells, hereafter M2NM8; 80% mobile and 20% non-mobile cells, hereafter M8NM2).

To compare the fitness of each population under standard culture conditions, supernatants of cultures of the growth experiment (day 10) were first collected by centrifugation at 5000 rpm for 2 min followed by syringe filtration (0.22 µm). Then, a starting density (∼1*10^4^ cells/mL) of each population was transferred into 100 mL of supernatant. Cell number was checked with a cell counting chamber under the microscope every 24 hours. The experiment lasted for 11 days and was done in triplicate.

For fitness assay under stressful conditions, salinity stress (extra 300 mM sodium chloride, NaCl, Karl Roth, Germany), oxidative stress (1 mM hydrogen peroxide, H_2_O_2_, Karl Roth, Germany) and drought stress were included. For salinity stress and oxidative stress, a starting density of ∼1*10^4^ cells/mL of each population was cultured in 200 mL of customed supernatant, respectively. For drought stress, 2 mL culture (∼1*10^4^ cells/mL) of each population was spread on a 0.22 µm membrane filter (round shape, 47 mm in diameter, Thermo Fisher Scientific, USA) upon a kitchen paper to remove the liquid. The filter membrane was then transferred into a petri dish and dried in the air. All triplicated treatments were exposed to stresses for 10 days, after which the surviving cells were enumerated under the microscope. Viable cells were discriminated from dead cells based on their intact cell shape and autofluorescence under the microscope. Unless otherwise specified, other culture parameters were 25±1°C under 12h12h light-dark cycles (20 µmol photons m^−2^ s^−1^).

### Re-diversification of the individual phenotypes

Triplicated mobile and non-mobile populations were each transferred for 10 days into 200 mL of fresh BBM medium at an initial density of 1*10^4^ cells/mL. The cell number and composition of the mobile population were checked daily.

### Statistical analyses

All statistical analyses were performed in R v4.1.1. Statistical significance for physiological measurements and nutrient dynamics was calculated by Welch’s *t* test for pairwise comparisons of two treatments (*p* value < 0.05). A one-way ANOVA with Tukey’s HSD post-hoc analysis (*p* value < 0.05) was conducted for all tested populations of growth and survival assays.

## Acknowledgements

This work was supported by the S&T Projects of Shenzhen Science and Technology Innovation Committee (JCYJ20200109142822787, KCXFZ2022, RCJC20200714114433069, and JCYJ20200109142818589), the Project of Shenzhen Municipal Bureau of Planning and Natural Resources (Grant No. [2021]735-927), as well as the Shenzhen-Hong Kong-Macao Joint S&T Project (pending number 202205303000176).

## Author contributions

S.T. and ZH.C. designed the experiments. S.T., Yaqing Liu and Xueyu Cheng performed the experiments. All authors interpreted the results and wrote the manuscript.

## Data availability

The growth parameters, growth and survival assay data and physiological data collected from this study are available from Zenodo online (https://doi.org/10.5281/zenodo.8328781). Raw RNAseq reads for differential gene expression analyses have been submitted to NCBI’s SRA database (http://www.ncbi.nlm.nih.gov) under BioProject PRJNA940855. All scripts for data visualization are available at https://github.com/SiTANG1990/Microalgal-bet-hedging.git.

---

**Figure. S1.**
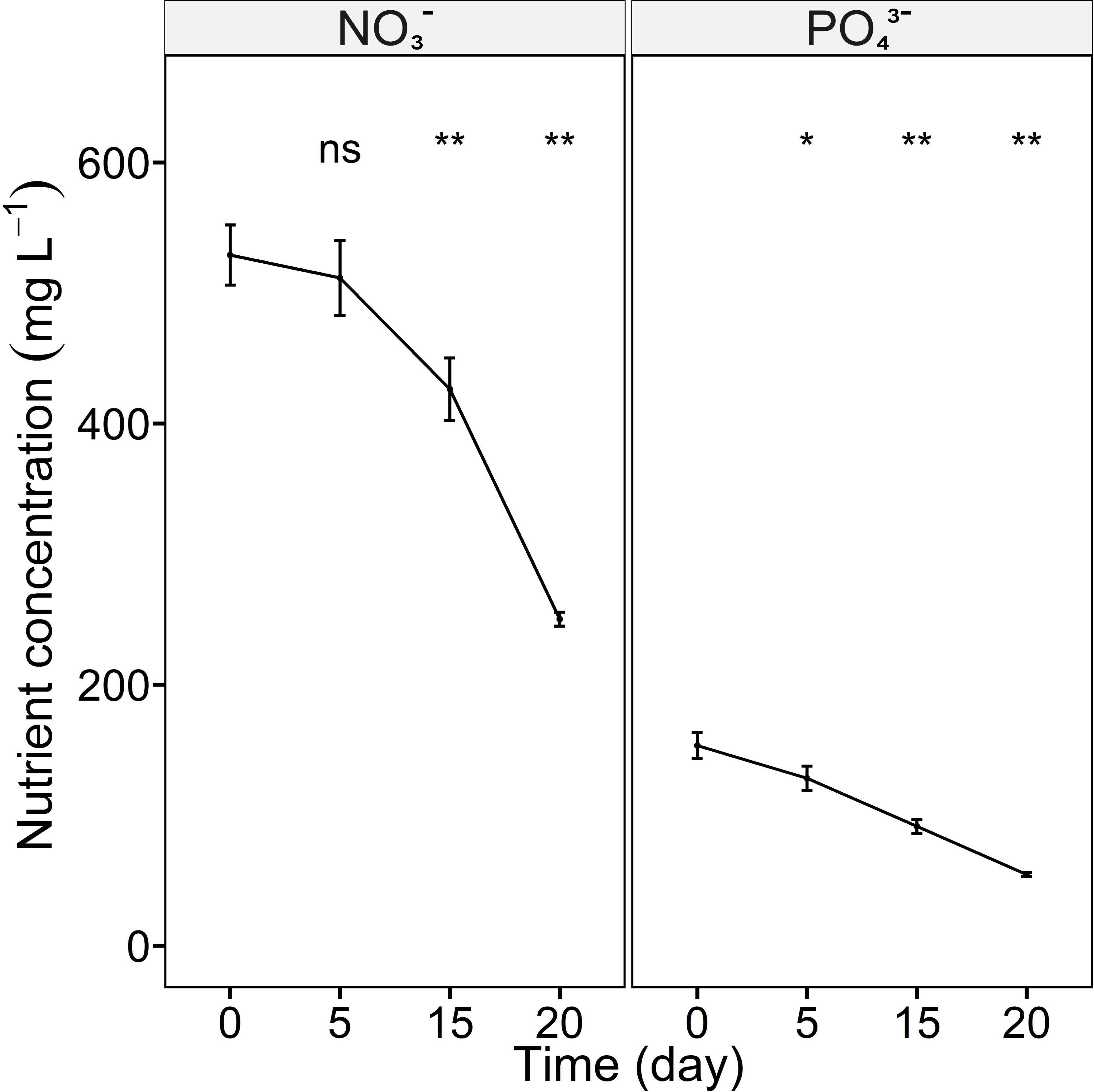
Nutrient dynamics over time. Dissolved NO_3-_ and PO_43-_ concentrations were quantified every five days. Error bars indicate the standard deviation of the triplicate values. Asterisks represent the statistical significance calculated by Welch’s *t* test between the data of the different time points and those of day 0. Significance: ns (no significance), *(*p* < 0.05), **(*p* < 0.01), ***(*p* < 0.001), ****(*p* < 0.0001).

## References

1. Lean, J. L. & Rind, D. H. How natural and anthropogenic influences alter global and regional surface temperatures: 1889 to 2006. Geophysical Research Letters 35, (2008).

2. Moran, M. A. et al. Deciphering ocean carbon in a changing world. Proceedings of the National Academy of Sciences 113, 3143–3151 (2016).

3. Philippi, T. & Seger, J. Hedging one’s evolutionary bets, revisited. Trends in Ecology & Evolution 4, 41–44 (1989).

4. Slatkin, M. Hedging one’s evolutionary bets. Nature 250, 704–705 (1974).

5. Veening, J.-W., Smits, W. K. & Kuipers, O. P. Bistability, Epigenetics, and Bet-Hedging in Bacteria. Annual Review of Microbiology 62, 193–210 (2008).

6. Veening, J.-W. et al. Bet-hedging and epigenetic inheritance in bacterial cell development. Proceedings of the National Academy of Sciences 105, 4393–4398 (2008).

7. Grimbergen, A. J., Siebring, J., Solopova, A. & Kuipers, O. P. Microbial bet-hedging: the power of being different. Current Opinion in Microbiology 25, 67–72 (2015).

8. Lycus, P. et al. A bet-hedging strategy for denitrifying bacteria curtails their release of N2O. Proceedings of the National Academy of Sciences 115, 11820–11825 (2018).

9. Ackermann, M. et al. Self-destructive cooperation mediated by phenotypic noise. Nature 454, 987–990 (2008).

10. Beaumont, H. J. E., Gallie, J., Kost, C., Ferguson, G. C. & Rainey, P. B. Experimental evolution of bet hedging. Nature 462, 90–93 (2009).

11. Childs, D. Z., Metcalf, C. J. E. & Rees, M. Evolutionary bet-hedging in the real world: empirical evidence and challenges revealed by plants. Proceedings of the Royal Society B: Biological Sciences 277, 3055–3064 (2010).

12. Danforth, B. N. Emergence dynamics and bet hedging in a desert bee, Perdita portalis. Proceedings of the Royal Society of London. Series B: Biological Sciences 266, 1985– 1994 (1999).

13. Levy, S. F., Ziv, N. & Siegal, M. L. Bet Hedging in Yeast by Heterogeneous, Age-Correlated Expression of a Stress Protectant. PLOS Biology 10, e1001325 (2012).

14. Olofsson, H., Ripa, J. & Jonzén, N. Bet-hedging as an evolutionary game: the tradeoff between egg size and number. Proceedings of the Royal Society B: Biological Sciences 276, 2963–2969 (2009).

15. Ratcliff, W. C. & Denison, R. F. Individual-Level Bet Hedging in the Bacterium Sinorhizobium meliloti. Current Biology 20, 1740–1744 (2010).

16. Venable, D. L. Bet Hedging in a Guild of Desert Annuals. Ecology 88, 1086–1090 (2007).

17. Arrigo, K. R. et al. Phytoplankton Community Structure and the Drawdown of Nutrients and CO2 in the Southern Ocean. Science 283, 365–367 (1999).

18. Field, C. B., Behrenfeld, M. J., Randerson, J. T. & Falkowski, P. Primary Production of the Biosphere: Integrating Terrestrial and Oceanic Components. Science 281, 237–240 (1998).

19. Li, W. K. W. Primary production of prochlorophytes, cyanobacteria, and eucaryotic ultraphytoplankton: Measurements from flow cytometric sorting. Limnology and Oceanography 39, 169–175 (1994).

20. Richardson, T. L. & Jackson, G. A. Small Phytoplankton and Carbon Export from the Surface Ocean. Science 315, 838–840 (2007).

21. Thomas, M. K., Kremer, C. T., Klausmeier, C. A. & Litchman, E. A Global Pattern of Thermal Adaptation in Marine Phytoplankton. Science 338, 1085–1088 (2012).

22. Chan, W. Y., Oakeshott, J. G., Buerger, P., Edwards, O. R. & van Oppen, M. J. H. Adaptive responses of free-living and symbiotic microalgae to simulated future ocean conditions. Global Change Biology 27, 1737–1754 (2021).

23. Boussiba, S. Carotenogenesis in the green alga Haematococcus pluvialis: Cellular physiology and stress response. Physiologia Plantarum 108, 111–117 (2000).

24. Khoo, K. S. et al. Recent advances in biorefinery of astaxanthin from Haematococcus pluvialis. Bioresource Technology 288, 121606 (2019).

25. Ren, Y. et al. Using green alga Haematococcus pluvialis for astaxanthin and lipid co-production: Advances and outlook. Bioresource Technology 340, 125736 (2021).

26. Shah, Md. M. R., Liang, Y., Cheng, J. J. & Daroch, M. Astaxanthin-Producing Green Microalga Haematococcus pluvialis: From Single Cell to High Value Commercial Products. Frontiers in Plant Science 7, (2016).

27. Orosa, M., Franqueira, D., Cid, A. & Abalde, J. Analysis and enhancement of astaxanthin accumulation in Haematococcus pluvialis. Bioresource Technology 96, 373–378 (2005).

28. Philippi, T. & Seger, J. Hedging one’s evolutionary bets, revisited. Trends in Ecology & Evolution 4, 41–44 (1989).

29. Starrfelt, J. & Kokko, H. Bet-hedging—a triple tradeoff between means, variances and correlations. Biological Reviews 87, 742–755 (2012).

30. Ashraf, M. & Harris, P. J. C. Photosynthesis under stressful environments: An overview. Photosynthetica 51, 163–190 (2013).

31. Ledford, H. K. & Niyogi, K. K. Singlet oxygen and photo-oxidative stress management in plants and algae. *Plant*, Cell & Environment 28, 1037–1045 (2005).

32. Vitova, M., Bisova, K., Kawano, S. & Zachleder, V. Accumulation of energy reserves in algae: From cell cycles to biotechnological applications. Biotechnology Advances 33, 1204–1218 (2015).

33. Zhang, L. et al. Salinity-induced cellular cross-talk in carbon partitioning reveals starch-to- lipid biosynthesis switching in low-starch freshwater algae. Bioresource Technology 250, 449–456 (2018).

34. Thalmann, M. & Santelia, D. Starch as a determinant of plant fitness under abiotic stress. New Phytologist 214, 943–951 (2017).

35. Pi, X. et al. Unique organization of photosystem I–light-harvesting supercomplex revealed by cryo-EM from a red alga. Proceedings of the National Academy of Sciences 115, 4423–4428 (2018).

36. Qin, X. et al. Structure of a green algal photosystem I in complex with a large number of light-harvesting complex I subunits. Nat. Plants 5, 263–272 (2019).

37. Cunningham, F. X. & Gantt, E. Genes and Enzymes of Carotenoid Biosynthesis in Plants. Annual Review of Plant Physiology and Plant Molecular Biology 49, 557–583 (1998).

38. Lohr, M., Im, C.-S. & Grossman, A. R. Genome-based examination of chlorophyll and carotenoid biosynthesis in Chlamydomonas reinhardtii. Plant Physiol 138, 490–515 (2005).

39. Lu, S. & Li, L. Carotenoid Metabolism: Biosynthesis, Regulation, and Beyond. Journal of Integrative Plant Biology 50, 778–785 (2008).

40. Bai, Y. et al. Green diatom mutants reveal an intricate biosynthetic pathway of fucoxanthin. Proceedings of the National Academy of Sciences 119, e2203708119 (2022).

41. Sampath, H. & Ntambi, J. M. Polyunsaturated Fatty Acid Regulation of Genes of Lipid Metabolism. Annual Review of Nutrition 25, 317–340 (2005).

42. Kazakov, A. E. et al. Comparative Genomics of Regulation of Fatty Acid and Branched- Chain Amino Acid Utilization in Proteobacteria. Journal of Bacteriology 191, 52–64 (2009).

43. Stitt, M. & Zeeman, S. C. Starch turnover: pathways, regulation and role in growth. Current Opinion in Plant Biology 15, 282–292 (2012).

44. Zhou, H. et al. Critical roles of soluble starch synthase SSIIIa and granule-bound starch synthase Waxy in synthesizing resistant starch in rice. Proceedings of the National Academy of Sciences 113, 12844–12849 (2016).

45. Delrue, B. et al. Waxy Chlamydomonas reinhardtii: monocellular algal mutants defective in amylose biosynthesis and granule-bound starch synthase activity accumulate a structurally modified amylopectin. Journal of Bacteriology 174, 3612–3620 (1992).

46. Ball, S. G. & Morell, M. K. From Bacterial Glycogen to Starch: Understanding the Biogenesis of the Plant Starch Granule. Annual Review of Plant Biology 54, 207–233 (2003).

47. Janeček, Š., Mareček, F., MacGregor, E. A. & Svensson, B. Starch-binding domains as CBM families–history, occurrence, structure, function and evolution. Biotechnology Advances 37, 107451 (2019).

48. Kim, H.-R. et al. Raw starch fermentation to ethanol by an industrial distiller’s yeast strain of Saccharomyces cerevisiae expressing glucoamylase and α-amylase genes. Biotechnol Lett 33, 1643–1648 (2011).

49. Viola, R., Nyvall, P. & Pedersén, M. The unique features of starch metabolism in red algae. Proceedings of the Royal Society of London. Series B: Biological Sciences 268, 1417–1422 (2001).

50. Zhang, S. & Klessig, D. F. MAPK cascades in plant defense signaling. Trends in Plant Science 6, 520–527 (2001).

51. Pitzschke, A., Schikora, A. & Hirt, H. MAPK cascade signalling networks in plant defence. Current Opinion in Plant Biology 12, 421–426 (2009).

52. Dóczi, R., Ökrész, L., Romero, A. E., Paccanaro, A. & Bögre, L. Exploring the evolutionary path of plant MAPK networks. Trends in Plant Science 17, 518–525 (2012).

53. Grimbergen, A. J., Siebring, J., Solopova, A. & Kuipers, O. P. Microbial bet-hedging: the power of being different. Current Opinion in Microbiology 25, 67–72 (2015).

54. Simões, M. et al. The Evolving Theory of Evolutionary Radiations. Trends in Ecology & Evolution 31, 27–34 (2016).

55. Hallet, B. Playing Dr Jekyll and Mr Hyde: combined mechanisms of phase variation in bacteria. Current Opinion in Microbiology 4, 570–581 (2001).

56. Darmon, E. & Leach, D. R. F. Bacterial Genome Instability. Microbiology and Molecular Biology Reviews 78, 1–39 (2014).

57. Bergmiller, T. & Ackermann, M. Pole Age Affects Cell Size and the Timing of Cell Division in *Methylobacterium extorquens* AM1. Journal of Bacteriology 193, 5216–5221 (2011).

58. Levy, S. F., Ziv, N. & Siegal, M. L. Bet Hedging in Yeast by Heterogeneous, Age- Correlated Expression of a Stress Protectant. PLOS Biology 10, e1001325 (2012).

59. Barrett, R. D. H. & Schluter, D. Adaptation from standing genetic variation. Trends in Ecology & Evolution 23, 38–44 (2008).

60. Bell, G. & Collins, S. Adaptation, extinction and global change. Evolutionary Applications 1, 3–16 (2008).

61. Beaumont, H. J. E., Gallie, J., Kost, C., Ferguson, G. C. & Rainey, P. B. Experimental evolution of bet hedging. Nature 462, 90–93 (2009).

62. Ratcliff, W. C. & Denison, R. F. Individual-Level Bet Hedging in the Bacterium *Sinorhizobium meliloti*. Current Biology 20, 1740–1744 (2010).

63. Solopova, A. et al. Bet-hedging during bacterial diauxic shift. Proceedings of the National Academy of Sciences 111, 7427–7432 (2014).

64. Lycus, P. et al. A bet-hedging strategy for denitrifying bacteria curtails their release of N2O. Proceedings of the National Academy of Sciences 115, 11820–11825 (2018).

65. Veening, J.-W., Smits, W. K. & Kuipers, O. P. Bistability, Epigenetics, and Bet-Hedging in Bacteria. Annual Review of Microbiology 62, 193–210 (2008).

66. Beaumont, H. J. E., Gallie, J., Kost, C., Ferguson, G. C. & Rainey, P. B. Experimental evolution of bet hedging. Nature 462, 90–93 (2009).

67. de Jong, I. G., Haccou, P. & Kuipers, O. P. Bet hedging or not? A guide to proper classification of microbial survival strategies. BioEssays 33, 215–223 (2011).

68. Ackermann, M. A functional perspective on phenotypic heterogeneity in microorganisms. Nat Rev Microbiol 13, 497–508 (2015).

69. Boussiba, S. Carotenogenesis in the green alga *Haematococcus pluvialis*: Cellular physiology and stress response. Physiologia Plantarum 108, 111–117 (2000).

70. Hagen, C., Siegmund, S. & Braune, W. Ultrastructural and chemical changes in the cell wall of *Haematococcus pluvialis* (Volvocales, Chlorophyta) during aplanospore formation. European Journal of Phycology 37, 217–226 (2002).

71. Thalmann, M. & Santelia, D. Starch as a determinant of plant fitness under abiotic stress. New Phytologist 214, 943–951 (2017).

72. Takaichi, S. Carotenoids in Algae: Distributions, Biosyntheses and Functions. Marine Drugs 9, 1101–1118 (2011).

73. Li-Beisson, Y., Thelen, J. J., Fedosejevs, E. & Harwood, J. L. The lipid biochemistry of eukaryotic algae. Progress in Lipid Research 74, 31–68 (2019).

74. Stahl, W. & Sies, H. Antioxidant activity of carotenoids. Molecular Aspects of Medicine 24, 345–351 (2003).

75. Christaki, E., Bonos, E., Giannenas, I. & Florou-Paneri, P. Functional properties of carotenoids originating from algae. Journal of the Science of Food and Agriculture 93, 5– 11 (2013).

76. A, K. T., et al. Cold-tolerant strain of *Haematococcus pluvialis* (Haematococcaceae, Chlorophyta) from Blomstrandhalvøya (Svalbard). ALGAE 28, 185–192 (2013).

77. Burchardt, L., Balcerkiewicz, S., Kokociński, M., Samardakiewicz, S. & Adamski, Z. Occurrence of *Haematococcus pluvialis* Flotow emend. Wille in a small artificial pool on the university campus of the Collegium Biologicum in Poznań (Poland). Biodiv. Res. Conserv. 2006, 163–166.

78. Czygan, F.-C. Blutregen und Blutschnee: Stickstoffmangel-Zellen von Haematococcus pluvialis und Chlamydomonas nivalis. Arch. Mikrobiol. 74, 69–76 (1970).

79. Kim, J. H. et al. Morphological, Molecular, and Biochemical Characterization of Astaxanthin-Producing Green Microalga Haematococcus sp. KORDI03 (Haematococcaceae, Chlorophyta) Isolated from Korea. Journal of Microbiology and Biotechnology 25, 238–246 (2015).

80. Xv, V., et al. Astaxanthin production from a new strain of *Haematococcus pluvialis* grown in batch culture. Annals of the Romanian Society for Cell Biology 15, (2010).

81. Evans, M. E. K. & Dennehy, J. J. Germ Banking: Bet-Hedging and Variable Release from Egg and Seed Dormancy. The Quarterly Review of Biology 80, 431–451 (2005).

82. Fábregas, J., Domínguez, A., Regueiro, M., Maseda, A. & Otero, A. Optimization of culture medium for the continuous cultivation of the microalga Haematococcus pluvialis. Appl Microbiol Biotechnol 53, 530–535 (2000).

83. Yan, Q. et al. Internal nutrient loading is a potential source of eutrophication in Shenzhen Bay, China. Ecological Indicators 127, 107736 (2021).

84. Simon, D. & Helliwell, S. Extraction and quantification of chlorophyll a from freshwater green algae. Water Research 32, 2220–2223 (1998).

85. Liu, Y. et al. A growth-boosting synergistic mechanism of Chromochloris zofingiensis under mixotrophy. Algal Research 66, 102812 (2022).

86. Saini, R. K. & Keum, Y.-S. Carotenoid extraction methods: A review of recent developments. Food Chemistry 240, 90–103 (2018).

87. Brányiková, I. et al. Microalgae—novel highly efficient starch producers. Biotechnology and Bioengineering 108, 766–776 (2011).

88. Chen, W., Zhang, C., Song, L., Sommerfeld, M. & Hu, Q. A high throughput Nile red method for quantitative measurement of neutral lipids in microalgae. Journal of Microbiological Methods 77, 41–47 (2009).

